# A Nickel N-Heterocyclic Biscarbene Complex Derived from Caffeine Enhances Fluconazole Efficacy against *Candida glabrata*

**DOI:** 10.64898/2026.03.03.709283

**Authors:** Cláudia Malta-Luís, Giulia Romeo, Giulia Francescato, Carolina Mariano, Dalila Mil-Homens, Ana Petronilho, Catarina Pimentel

**Affiliations:** Instituto de Tecnologia Química e Biológica António Xavier, Universidade Nova de Lisboa, Av. República, 2780-157 Oeiras, Portugal

**Author notes:** Adress correspondence to Ana Petronilho, and Catarina Pimentel,.

## Abstract

Invasive infections caused by *Candida* spp. are associated with high morbidity and mortality, particularly in immunocompromised patients, and are increasingly difficult to treat due to rising antifungal resistance. Here, we further investigate the antifungal properties of a previously reported nickel N-heterocyclic biscarbene derived from caffeine (compound 5) and describe the synthesis of an analogous nickel N-heterocyclic biscarbene based on benzimidazole (compound 8), designed to evaluate the impact of the biscarbene and xanthine frameworks on activity and toxicity. Compound 5 displayed selective activity against *Candida glabrata*, inhibited biofilm formation and showed greater cellular accumulation in this species. In a *Galleria mellonella* infection model, 5 significantly reduced fungal burden while exhibiting lower cytotoxicity than the benzimidazole analogue 8. Importantly, although compound 5 and fluconazole are individually fungistatic, their combination was fungicidal against *C. glabrata*. In the presence of compound 5, the minimal inhibitory concentration of fluconazole decreased against both a fluconazole-resistant *petite* mutant and a respiratory-competent *C. glabrata* strain evolved *in vitro* to fluconazole resistance. Compound 5 increased the frequency of *petite* mutants, suggesting an effect on mitochondrial function; however, its retained activity against these mutants and its synergism with fluconazole in the *petite* background indicate an additional mechanism of action. These findings identify 5 as a promising antifungal adjuvant and support its potential use in combination therapy to enhance azole efficacy against *C. glabrata*.

**Importance:** Invasive infections caused by *C. glabrata* are increasingly hard to treat because this pathogen is often tolerant to azoles and is developing resistance to echinocandins, leaving clinicians with few effective options. Our work explores a nickel–caffeine complex that shows selective activity against *C. glabrata* and low toxicity, features that make it attractive as a potential therapeutic lead. Notably, this compound enhances the activity of fluconazole, turning a commonly used fungistatic drug into a fungicidal combination against both susceptible and resistant *C. glabrata*. By also affecting mitochondrial function yet remaining active in respiratory-deficient mutants, the compound appears to act through mechanisms distinct from existing antifungals. These properties suggest that metal-based xanthine complexes could be developed as adjuvants to restore or boost azole efficacy against difficult-to-treat *C. glabrata* infections, addressing an important priority identified for fungal pathogens.

## Introduction

Invasive fungal infections caused by *Candida* spp are associated with high morbidity and mortality worldwide, especially among immunocompromised patients (1). To manage these infections, only a few antifungal classes, namely polyenes, azoles and echinocandins, are available, however, the number of clinical strains resistant to them has increased over the years, resulting in infections that are difficult or sometimes impossible to treat (2, 3). Prompted by this scenario, the World Health Organization has published the first-ever fungal priority list, categorizing some *Candida* species into the “critical priority group”, and highlighting the urgent need for novel antifungals with distinct mechanisms of action (4, 5).

Although *Candida albicans* remains the most common cause of invasive candidiasis, non-*Candida albicans* species are emerging as clinical challenges (1). Depending on the geographic area, *Candida glabrata* (currently classified as *Nakaseomyces glabrata* (6)) is the second most frequent cause of invasive candidiasis in hospital settings. To overcome the intrinsically low susceptibility of *C. glabrata* to azoles (6–8), echinocandins are typically recommended as first-line treatment for invasive candidiasis caused by this species. Nevertheless, the incidence of *C. glabrata* clinical isolates resistant to this class of antifungals is also increasing, with approximately 14% of fluconazole-resistant isolates exhibiting resistance to at least one echinocandin (9–11).

Metal-based compounds have re-emerged as promising antifungal agents in response to current treatment challenges (12, 13). These compounds show stronger antifungal activity than organic molecules because of physicochemical properties uniquely conferred by the metals (12–14). In particular, nickel(II) has attracted attention because, in addition to its abundance and low cost compared to other transition metals, it exhibits versatile coordination chemistry and redox behavior, which allow the formation of diverse structures with multiple mechanisms of action (13, 15–17).

We have previously reported the synthesis and antifungal activity of several nickel complexes bearing N-heterocyclic carbene precursors (NHCs) derived from xanthines (21). Xanthines have been widely used as ligands in metal complexes, since they offer to the structure electronic flexibility, strong donor ability and compatibility with a wide range of transition metals (18–20). Within that series, it was particularly noteworthy that the only biscarbene derivative (compound **5**), based on caffeine, exhibited remarkably low *in vitro* toxicity compared to its monocarbene counterpart, while maintaining selectivity against *C. glabrata*.

In this work, the effect of introducing an additional carbene was examined through the synthesis of a different compound, benzimidazole-based N-heterocyclic biscarbene (compound **8)**, demonstrating that the presence of an extra carbene moiety alone is insufficient to reduce cytotoxicity. Although compound **8** exhibited greater *in vitro* activity than compound **5**, this trend was not observed *in vivo*, where compound **5** showed superior efficacy in a *Galleria mellonella* model of invasive candidiasis caused by *C. glabrata*. The selectivity and antifungal activity of compound **5**, together with its synergistic interaction with the most widely used azole, fluconazole, were further evaluated, highlighting its potential as an adjuvant in combination therapy for overcoming antifungal resistance.

## Results

### Synthesis of compound 8

The benzimidazium salt I (22) was reacted with nickelocene at 100°C overnight and after appropriate work-up, compound **8** was obtained in 22% yield (**Scheme 1**). Analysis of the solid by ^1^H NMR confirms the disappearance of H1 and the appearance of diagnostic multiplets at 7.60 and 7.30 ppm (Fig. S1). In ^13^C the carbenic carbon resonates at 178.6 ppm (Fig. S2), confirmed by 2D correlation experiments, a value similar to those of similar bis-NHCs reported for nickel (23).

**Scheme 1.**
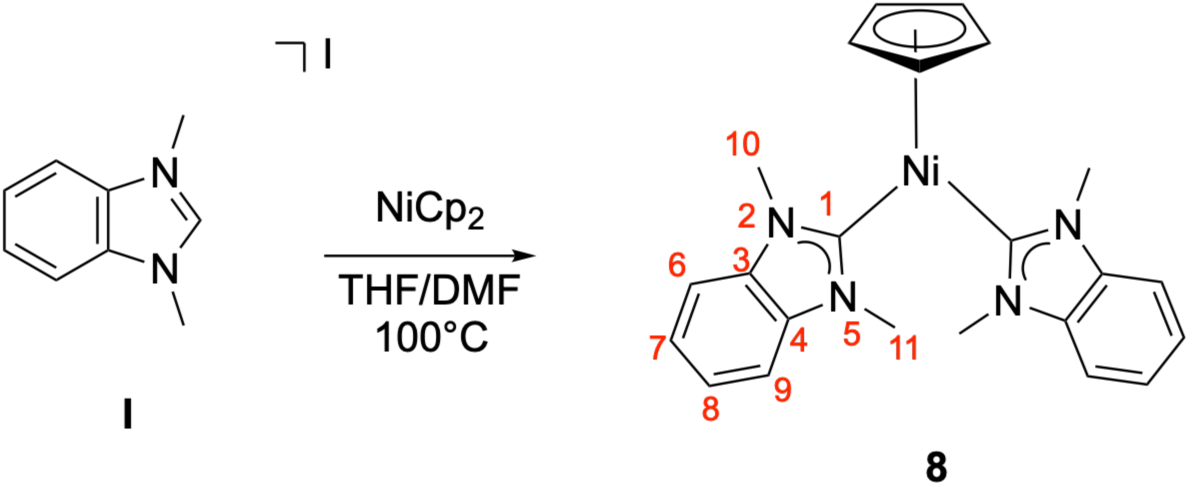
Synthesis of compound 8.

Compound **8** is characterized by a similar structure than that of compound **5** (Fig. 1), with both complexes being bis-NHCs based on fused heterocyclic rings. Their distinct nature stems from the type of N-heterocycle involved as ligand. Compound **5** is based on the natural compound caffeine, a xanthine, for which biological relevance and role in medicinal applications is well documented.

**Figure 1.**
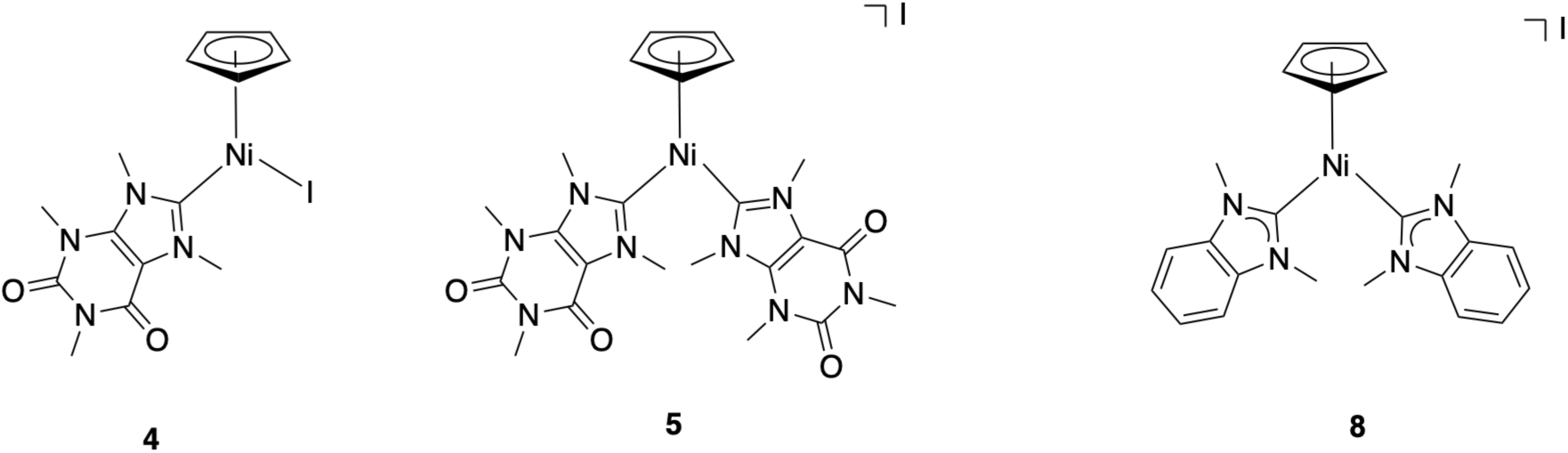
Molecular structure of bis-NHC complexes **5** and **8.**

### Effect of nickel complexes bearing N-heterocyclic biscarbene on antifungal activity and cytotoxicity

In a previous study, we showed that a biscarbene nickel complex of caffeine (**5**), compared to a monocarbene complex, not only increased the specificity of the compound toward *C. glabrata* but also reduced its cytotoxicity in HeLa cells (21). Therefore, we assessed whether the same effect could be observed with the newly synthesized nickel complex of an N-heterocyclic biscarbene from a different chemical family, benzimidazole (Fig. 1). The minimal inhibitory concentration (MIC) of **5** and **8** against the two most prevalent *Candida* spp associated with invasive candidiasis, *C. albicans* and *C. glabrata*, was determined according to the CLSI guidelines for yeast (24). As previously shown, compound **5** is highly selective for *C. glabrata*, exhibiting MICs 16-fold lower than those for *C. albicans* (Table 2 and (21)). Compound **8** is more effective against both species (lower MICs) than **5** (Table 2) or its monocarbene counterpart (75 μM for *C. glabrata* and 156 μM for *C. albicans*; compound **4** in Fig.1 (21)), but selectivity for *C. glabrata* was not observed.

**Table 1.** Yeast species used in this study.

|  | Origin |
| --- | --- |
| *Candida albicans* (SC5314) | ATCC |
| *Candida glabrata* (ATCC 2001) | ATCC |
| *Candida krusei* (ATCC6258) | ATCC |
| *Candida tropicalis* (ATCC 750) | ATCC |
| *Candida parapsilosis* (ATCC 22019) | ATCC |
| *Candida glabrata* resistant to fluconazole | (22) |
| *Saccharomyces cerevisiae* BY4742 | Euroscarf |

**Table 2.** Minimal inhibitory concentration (MIC) of 5 and 8 against *Candida albicans* and *Candida glabrata*.

|  | MIC (μM) |  |
| --- | --- | --- |
|  | 5 | 8 |
| *Candida albicans* | 1250 | 4.9 |
| *Candida glabrata* | 78.1 | 2.4 |

Despite its potent *in vitro* activity, compound **8** showed reduced efficacy *in vivo* in the *Galleria mellonella* model compared with compound **5** (Fig. 2A). Administration of 312.5 µM of compound **8** following *C. glabrata* infection did not reduce fungal burden in the larvae hemolymph, whereas treatment with 312.5 µM of compound **5** led to a significant decrease. In addition, compound **8** remained highly toxic to HeLa cells (Fig. 2B). Based on its promising *in vivo* activity and lower toxicity, compound **5** was selected for further biological characterization.

**Figure 2.**
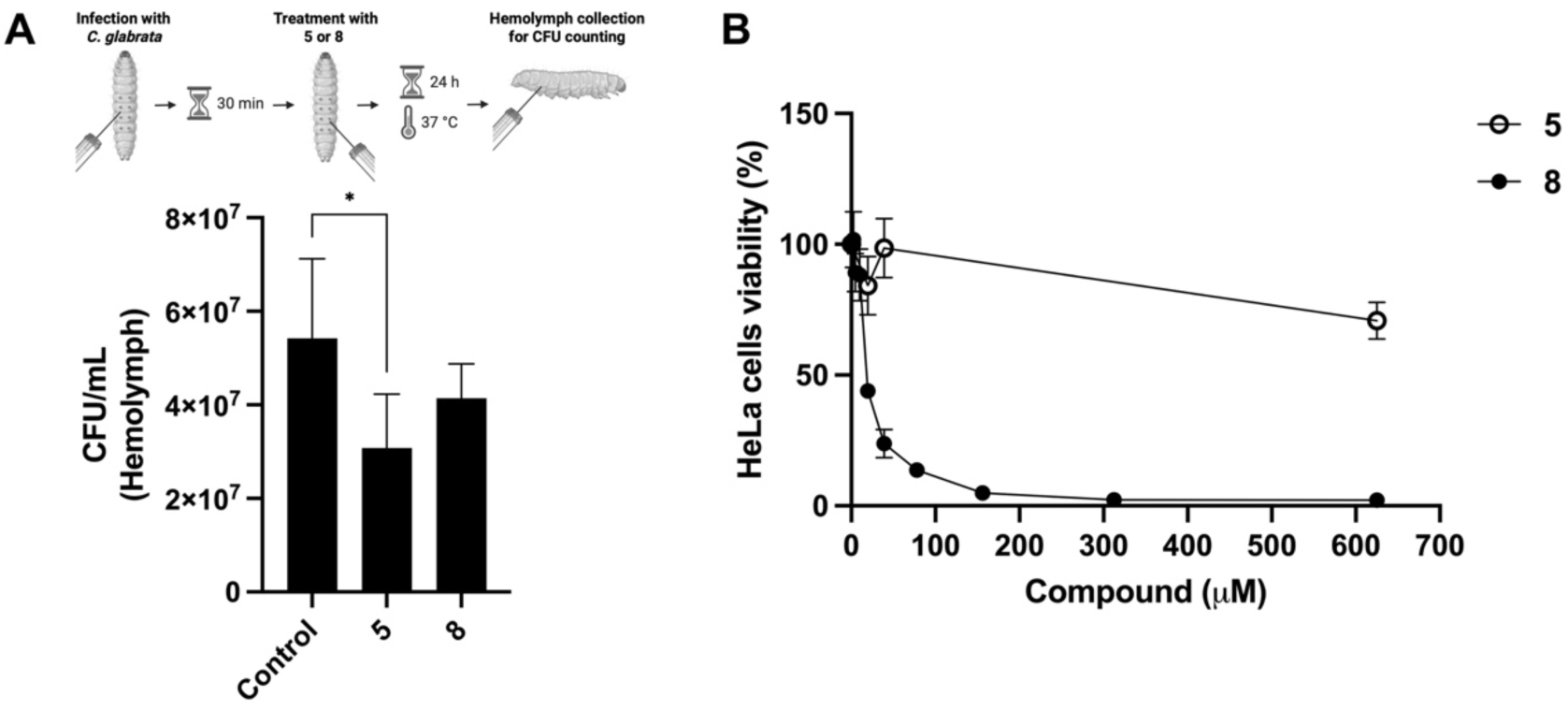
Compound 8 shows reduced *in vivo* activity against *C. glabrata* and higher cytotoxicity compared with compound 5. (A) Fungal burden in the hemolymph of infected *G. mellonella* treated with PBS (Control), 312.5 μM of compound **5** (**5**) or 312.5 μM of compound **8** (**8**). Hemolymph was collected from infected larvae after 24 h and plated on YPD agar plates for CFU counting. Statistical significance was assessed using one-way ANOVA with Turkey’s HSD post hoc test (*p≤0.05). (B) Cellular viability was evaluated using the MTT assay and the values are expressed as the percentage of viability relative to the untreated control.

The caffeine-based N-heterocyclic biscarbene 5 accumulates to higher levels in *C. glabrata* than in *C. albicans*.

To investigate the specificity of **5** towards *C. glabrata*, we quantified the nickel content of *C. albicans* and *C. glabrata* cells exposed to either 78.1 μM or 312.5 μM of **5**, (corresponding to the MIC and 4 x MIC for *C. glabrata*, respectively) using ICP-AES. At both concentrations, **5** accumulated to higher intracellular levels in *C. glabrata* than in *C. albicans* cells (Fig. 3), suggesting that its higher activity against *C. glabrata* may be linked to enhanced cellular uptake in this species.

**Figure 3.**
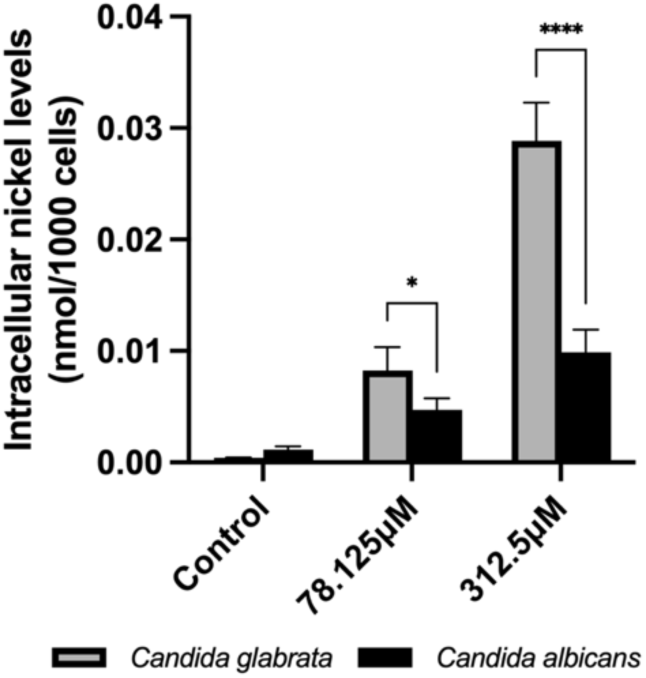
Higher intracellular levels of nickel after treatment with 5 were observed in *Candida glabrata*. The nickel content of *C. glabrata* cells, either untreated (Control) or treated with the indicated concentrations of 5, was quantified by ICP-AES. Data represent the mean ± standard deviation of four biological replicates. Statistical significance was assessed using two-way ANOVA with Turkey’s HSD post hoc test (*p≤0.05, ****p≤0.0001).

The spectrum of activity of compound **5** was also evaluated against other clinically relevant *Candida* spp included in the WHO fungal priority pathogens list, as well as against the baker’s yeast *Saccharomyces cerevisiae* (5). Consistent with its marked selectivity, **5** displayed activity against the phylogenetically related yeasts *C. glabrata* and *S. cerevisiae* and showed poor or no activity against other *Candida* spp within the tested concentration range (Table 3). Importantly, **5** was equally effective against a fluconazole-resistant *C. glabrata* strain.

**Table 3.** Minimal inhibitory concentration (MIC) of 5 against medically relevant *Candida* spp.

|  | MIC (μM) |
| --- | --- |
| *Candida albicans* | 1250 |
| *Candida glabrata* | 78.1 |
| *Candida krusei* | > 2500 |
| *Candida tropicalis* | > 2500 |
| *Candida parapsilosis* | > 2500 |
| *Candida glabrata* resistant to fluconazole* | 78.1 |
| *Saccharomyces cerevisiae* | 2.4 |

\*MIC<sub>fluc</sub> = 209 μM.

### Compound 5 acts synergistically with the antifungal fluconazole

A current trend in antifungal therapy research is the combination of antifungal agents with new or existing compounds that can act synergistically to improve antifungal potency and reduce the risk of resistance (25–27). Within this context, the possible synergistic effect of **5** with fluconazole, one of the most prescribed antifungal drugs worldwide, was tested. Using a checkerboard assay, we evaluated the growth of a wild-type *C. glabrata* strain (Fig. 4A) and a laboratory-evolved fluconazole-resistant *C. glabrata* strain (Fig. 4B) in response to a combination of two-fold serial dilutions of **5** (0.6-625 μM) and fluconazole (6.5-417.9 μM). After 24 h of growth, the wild-type strain showed a marked decrease in cell density when treated with a combination of compound **5** (9.8–39 μM) and fluconazole (6.5 μM), compared with equivalent concentrations of each drug alone (Fig. 4A, left panel). This effect was further supported by reduced cell viability, as assessed by spotting 5 μL of checkerboard assay cultures onto YPD agar plates (Fig. 4A, right panel). A similar synergistic pattern was also observed in the fluconazole-resistant strain (Fig. 4B).

**Figure 4.**
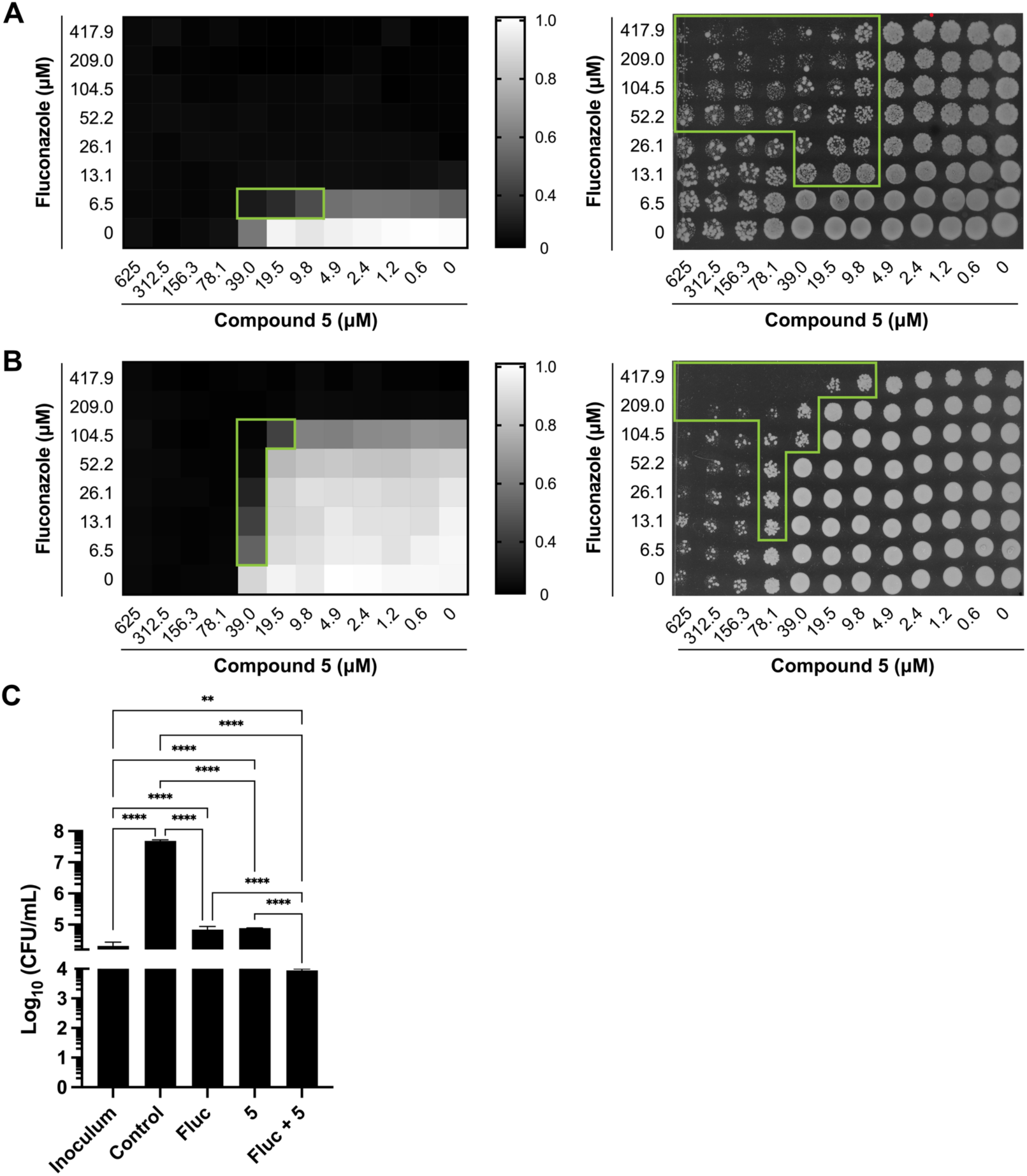
Synergistic interaction between compound 5 and fluconazole. Checkerboard assays were performed to evaluate the growth of (A) *C. glabrata* wild-type strain and (B) a fluconazole-resistant *C. glabrata* strain. After 24 h, OD_600_ was measured, values were normalized to untreated controls (left panels) and 5 μL of each culture was spotted onto YPD agar plates (right panels). (C) *C. glabrata* cell viability was further assessed by CFU counting of the initial inoculum (Inoculum), after 24h of growth (Control), or following treatment with 78.1 μM of **5** (5), 52.2 μM of fluconazole (Fluc), or their combination (Fluc + **5**). Results are expressed as Log_10_(CFU/mL) and represent the mean ± standard deviation of three biological replicates. Statistical differences were determined using one-way ANOVA on Log_10_-transformed CFU/mL values (**p≤0.01, ****p≤0.0001). The regions corresponding to concentrations at which the combination is more effective than the individual drugs alone are delineated by the green line.

Most importantly, although **5** (78.1 μM) and fluconazole (52.2 μM) exhibited fungistatic activity, reducing growth relative to the no-drug control without affecting cell viability compared with the initial inoculum, their combined treatment was fungicidal, as determined by colony-forming unit (CFU) counting (Fig. 4C).

To elucidate this synergistic effect, intracellular nickel levels were quantified by ICP-AES in *C. glabrata* cells co-treated with **5** and fluconazole. The combination resulted in a further increase in intracellular nickel content (Fig. 5A), suggesting that the observed synergy likely arises from enhanced cellular uptake of **5**, possibly due to fluconazole-induced alterations in sterol levels (28).

**Figure 5.**
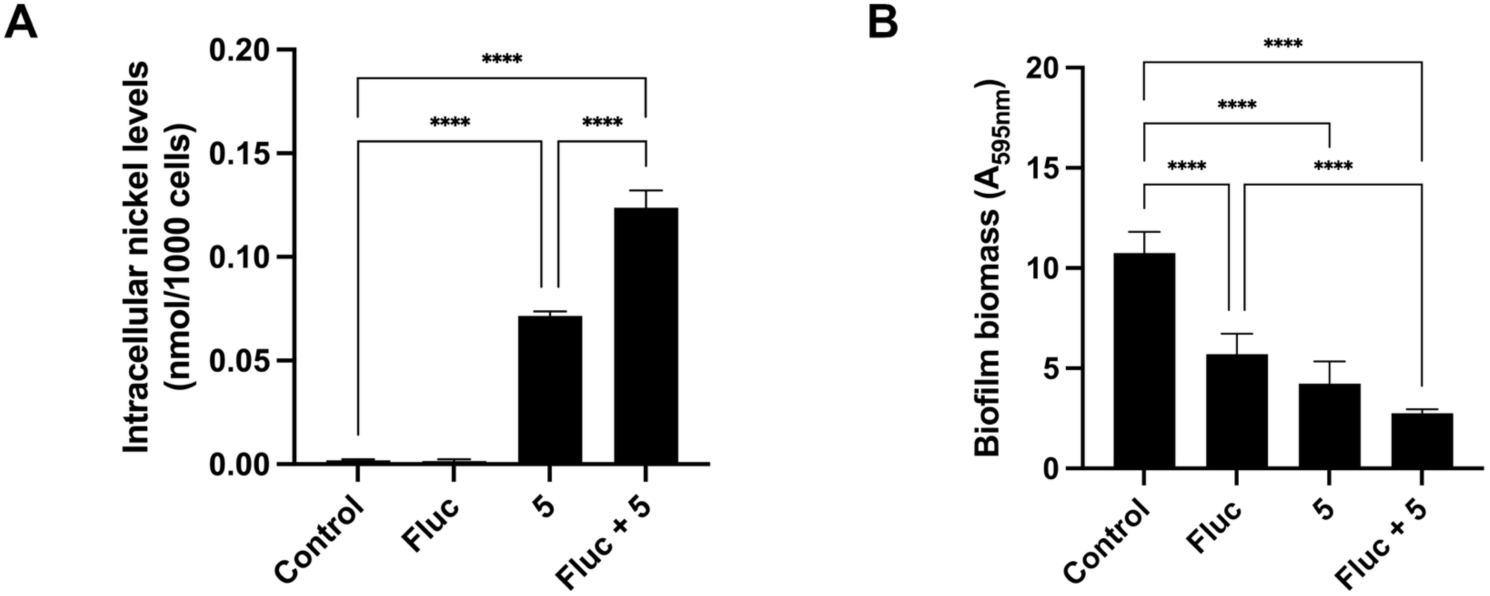
Combining compound 5 with fluconazole increases intracellular nickel levels and enhances biofilm inhibition. (A) Intracellular nickel content of *C. glabrata* cells, either untreated (Control) or treated with 78.1 μM of compound **5** (5), 52.2 μM of fluconazole (Fluc) or their combination (Fluc + 5), was quantified by ICP-AES. Data represent the mean ± standard deviation of four biological replicates. Statistical significance was assessed using one-way ANOVA with Turkey’s HSD post hoc test (****p≤0.0001). (B) Biofilm formation of *C. glabrata* either left untreated (Control) or treated with 78.1 μM of compound **5** (5), 52.2 μM of fluconazole (Fluc) or the corresponding concentrations of both (Fluc + 5). Biofilm mass was quantified using the crystal violet staining method. Values represent the mean of eight replicates. Statistical significance was assessed using one-way ANOVA with Turkey’s HSD post hoc test (**p≤0.01, ****p≤0.0001).

One key virulence factor of *Candida* spp is their ability to form biofilms on biotic and abiotic surfaces, contributing to persistent infections and increasing the risk of resistance development (29). Because caffeine has been reported to inhibit the formation of *C. albicans* biofilms (30), the impact of **5**, a caffeine derivative, alone and in combination with fluconazole, on *C. glabrata* biofilm formation was evaluated using the crystal violet staining method. Compound **5** strongly impaired biofilm development, and this effect was even more pronounced when combined with fluconazole (Fig. 5B).

### Treatment of *Candida glabrata* with compound 5 induces *petite* mutants

During cell viability assays (Fig. 4), we observed that treatment with **5**, either alone or in combination with fluconazole, led to the appearance of a subpopulation of slow-growing colonies, which was particularly striking under co-treatment conditions. These colonies were smaller and became visible to the naked eye only after an additional 24 h of incubation, consistent with the characteristics of *petite* mutants, which arise following partial or complete loss of mitochondrial function in yeast (31, 32).

Since *petite* mutants are respiratory-deficient and cannot grow on non-fermentable carbon sources (33), we randomly select slow-growing, small colonies and subculture them on YPD and YPG agar plates containing glucose or glycerol, respectively (Fig 6A). The inability of some colonies to grow on YPG confirmed their respiratory-deficient, *petite* nature. The formation of *petite* mutants was quantified by plating equal volumes of *C. glabrata* cultures, treated for 24 h with 78.1 μM of compound **5**, 52.2 μM of fluconazole, or their combination, onto YPD and YPG agar plates. CFUs were counted after 24h and 48h and the difference between YPD and YPG colony numbers was used to determine the number of *petite* mutants. While no *petite* mutants were observed during the growth period with fluconazole alone, approximately 70% of colonies exhibited respiratory deficiencies following treatment with compound 5, and 100% displayed this phenotype after co-treatment with both compounds (Fig. 6B).

**Figure 6.**
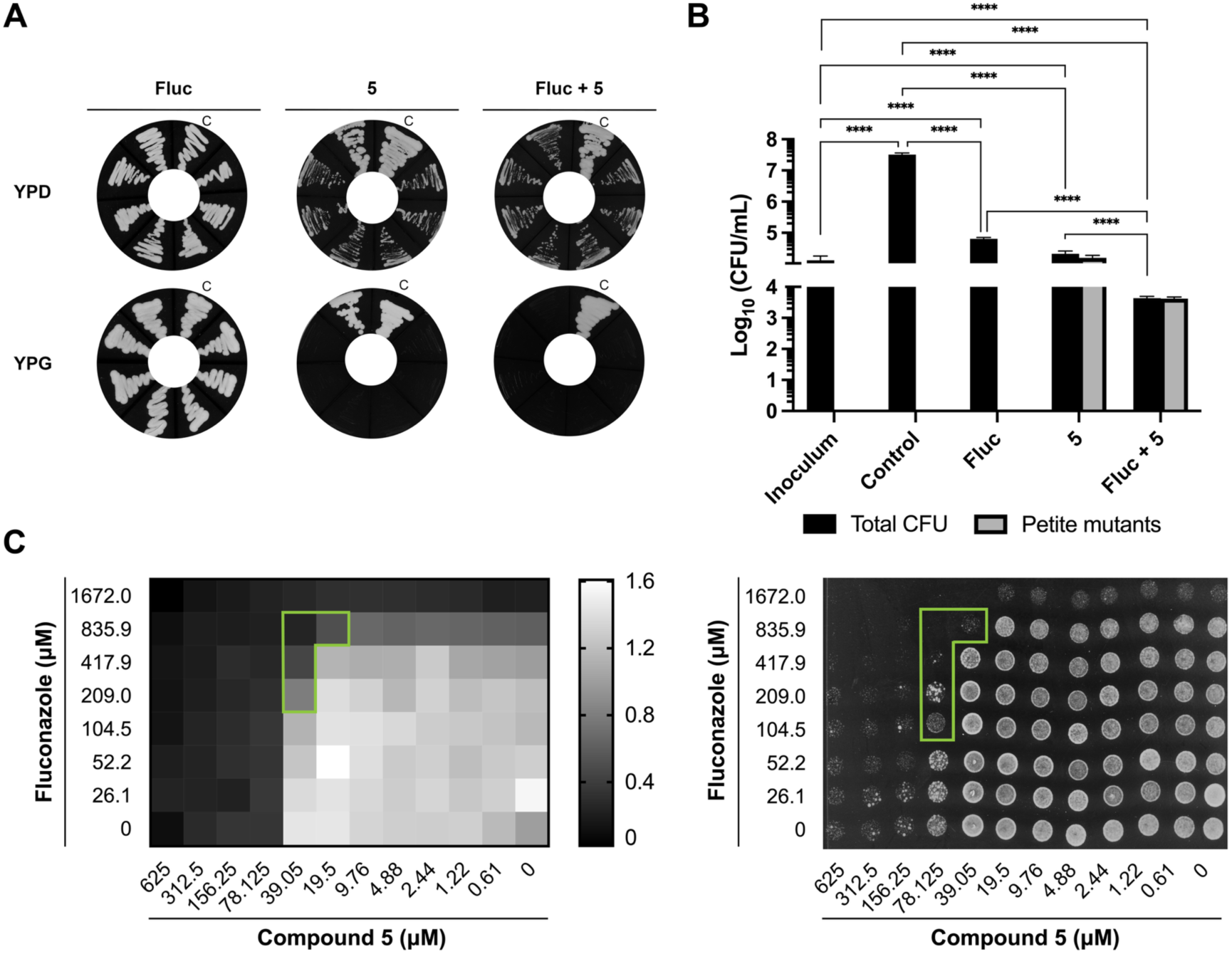
Compound 5 induces *petite* mutants. (A) Representative growth of slow-growing, small colonies on YPD and YPG agar plates. C-normal growing colony (B) Quantification of petite mutants. *C. glabrata* cultures were treated for 24 h with 78.1 μM compound 5 (5), 52.2 μM fluconazole (Fluc), or their combination (Fluc + 5). Statistical differences were determined using one-way ANOVA on Log_10_-transformed CFU/mL values (****p≤0.0001). (C) Checkerboard assays using a compound 5-induced petite mutant strain. After 24 h, OD_600_ was measured and values were normalized to untreated controls (left panel), and 5 μL of each culture was spotted onto YPD agar plates (right panel). The regions corresponding to concentrations at which the combination is more effective than the individual drugs alone are delineated by the green line.

*Petite* mutants are well known to be resistant to fluconazole (34), and those induced by compound **5** exhibit an MIC of 1672 μM, which is approximately 128-fold higher than that of the wild-type strain. We therefore questioned whether the synergy observed between compound **5** and fluconazole would be maintained in *petite* mutants. Checkerboard analysis using a compound **5**–induced *petite* mutant demonstrated that the synergistic effect remained evident (Fig. 6C), indicating that the interaction is not abolished by the *petite* phenotype.

## Discussion

The increasing number of *Candida* clinical isolates resistant to one or more antifungal agents, together with the limited pipeline of new antifungals, contributes to rising mortality rates, particularly among critically ill patients, underscoring the urgent need for novel antifungal therapies (**1, 2, 35**). In response to this clinical challenge, metal-based compounds have emerged as promising antifungal candidates (12, 13). Among transition metals, nickel has been highlighted in several reports as a metal of interest (13, 20, 21, 36–40). Structurally, most nickel(II) complexes described to date are Schiff-base ligands, leaving N-heterocyclic carbene-based nickel complexes largely underexplored.

We have previously identified compound **5**, a nickel(II) N-heterocyclic biscarbene complex, as exhibiting selective activity against *C. glabrata* and low cytotoxicity (21). Because the corresponding monocarbene was highly toxic to mammalian cells and less effective, we hypothesized that the biscarbene framework might confer an advantage. However, this was not the case for a nickel(II) biscarbene of benzimidazole (compound **8**). Although **8** showed greater *in vitro* antifungal activity than **5** and its monocarbene counterpart (21), it also exhibited higher *in vitro* toxicity. Despite being more lipophilic than **5** (logP 6.7 versus 1.4-theoretical calculation using MarvinSketch from Chemaxon™), **8** accumulated to lower intracellular levels (Fig. S3), suggesting potential sequestration within the plasma membrane (41). The higher lipophilic character of **8** may as well explain its low *in vivo* activity. In fact, compound efficacy in *Galleria mellonella* often does not correlate directly with *in vitro* potency. In a study of fenarimol analogues, compounds with lower lipophilicity were significantly more effective in increasing larval survival, whereas more lipophilic analogues showed reduced efficacy despite comparable *in vitro* activity, suggesting that high lipophilicity negatively affects *in vivo* performance in this model (42).

Compound **5** was selective for *C. glabrata* and *S. cerevisiae* but showed limited activity against other yeast species. *C. glabrata* is more closely related to *S. cerevisiae* than to other *Candida spp*, sharing a common ancestor that underwent whole genome duplication (43, 44). Therefore, a possible explanation is that uptake of **5** is mediated by a transporter that is specific to these species, exhibits higher affinity for **5**, or is overexpressed in these yeasts.

An interesting finding of this study is the induction of respiratory-deficient (*petite*) mutants by compound **5**. *Petite* mutants typically appear when cells have undergone total or partial loss of mitochondrial DNA (mtDNA) (32, 33), often caused by oxidative stress (45). Xanthine-derived NHC complexes have mainly been studied for their anticancer activity, and some reports suggest oxidative stress-related mechanisms for selected complexes (46–48). However, under our experimental conditions, we did not detect any increase in reactive oxygen species (ROS) after exposing *C. glabrata* to **5** (Fig. S4), making ROS induction an unlikely mechanism for this compound. Another possible route for *petite* formation, observed with DNA intercalating agents such as ethidium bromide, is the binding of the drug to mtDNA, which leads to the inhibition of replication and the induction of excessive DNA repair, ultimately resulting in mtDNA deletions (49). In *S. cerevisiae*, caffeine has been shown to be a *petite*-inducing agent (50, 51). Wolf *et al*. (50) proposed that this phenotype may arise from binding of caffeine to mtDNA, either by stacking externally on terminal purine bases or by intercalating between purine bases (52), leading to mispairing and ultimately to loss of mtDNA. In this context, since **5** is a caffeine derivative, a similar mechanism cannot be excluded.

The fact that **5** induces *petite* mutants was surprising in the context of its synergism with fluconazole, as in *C. glabrata*, *petite* mutants are well documented to exhibit reduced susceptibility to this azole fluconazole (34, 53, 54). In such cases, resistance has been attributed to mitochondrial dysfunction–driven retrograde signalling, which results in the constitutive upregulation of ATP-binding cassette (ABC) drug efflux pumps such as Cdr1 and Cdr2 (55, 56). One possible explanation for the observed synergism is that additional mutations or cellular alterations induced by compound **5** contribute to its antifungal activity. Further support for this hypothesis comes from two observations: (i) the synergism between fluconazole and **5** remains evident in *petite* mutants, and (ii) the proportion of *petite* cells is markedly reduced upon co-treatment with fluconazole compared to treatment with compound **5** alone. Together, these findings reconcile the apparent paradox whereby compound **5** induces fluconazole-resistant *petite* mutants yet acts synergistically with fluconazole. A similar mechanism may apply to caffeine, as it has been reported to reduce the proportion of *petite* mutants when combined with intercalating agents such as ethidium bromide (57, 58).

Although fluconazole did not induce *petite* mutants under our treatment conditions, it is well known to promote this phenotype *in vitro*, and *petite* clinical strains have been isolated from fluconazole-treated patients (34, 59, 60). In such cases, compound **5** may provide a promising co-treatment strategy to counteract the resistance mechanism triggered by these mutants and thereby enhance the therapeutic efficacy of fluconazole.

## Material and Methods

### Synthesis of compounds 5 and 8

The synthesis of compound **5** was previously described in (21). Synthesis of compound **8**: A Schlenk tube was charged with dimethylbenzimidazole (50 mg, 0.18 mmol) and nickelocene (34 mg, 0.18). After the addition of DMF (1.8 ml), the reaction was heated at 100 °C and stirred overnight. After cooling, the solvent was concentrated under vacuum and then Et_2_O was added to induce precipitation. The solid was separated by filtration, washed with Et_2_O and dried, yielding the desired compound as a brown solid in 22% yield (11 mg). ^1^H NMR (400 MHz, DMSO): δ ^1^H NMR (400 MHz, DMSO): δ 7.61-7.58 (m, 4H, H7-H8 Ar, ^3^J_H7-H6_ = ^3^J_H8-H9_ ≈ 6.0 Hz, ^4^J_H7-H9_ = ^4^J_H8-H6_ ≈ 3.1 Hz); 7.31-7.28 (m, 4H, H6-H9 Ar, ^3^J_H6-H7_ = ^3^J_H9-H8_ ≈ 6.0 Hz, ^4^J_H6-H8_ = ^4^J_H9-H7_ ≈ 3.1 Hz); 5.71 (s, 5H, C_5_H_5_); 4.23 (s, 12H, N2-CH_3_, N5-CH_3_) ppm.^13^C {^1^H} NMR (100 MHz, DMSO): δ 178.6 (C-1); 135.3 (C-3, C-4); 122.9 (C-6, C-9); 110.5 (C-7, C-8); 92.4 (C_5_H_5_); 35.8 (N2-CH_3_, N5-CH_3_) ppm. HRMS-ESI (m/z): calcd: 415.1433; found: 415.1420.

### Strains and growth conditions

The *Candida* spp used in this study are listed in Table 1. Strains were maintained in Sabouraud Dextrose Agar (SDA) plates. Unless otherwise indicated, all assays were performed in Roswell Park Memorial Institute 1640 (RPMI-1640) medium with 0.165 M of MOPS adjusted to pH 7.0. For ICP-AES assays, strains were grown in SC medium pH 5.5 (0.77 g/L complete supplement mixture; 1.7 g/L yeast nitrogen base w/o amino acids and ammonium sulphate; 5.4 g/L ammonium sulphate; 2% glucose). The wild-type (*C. glabrata* ATCC) and *petite* mutant strains were maintained in YPD medium (1% yeast extract, 2% peptone, 2% glucose and 2% agar). Cell viability was determined by counting colony-forming units (CFU). To this end, cultures were serially diluted in PBS and seeded onto YPD agar plates. The presence of *petite* mutants was examined on YPG agar plates (1% yeast extract, 2% peptone, 2% glycerol and 2% agar) and growth was recorded after 72h of incubation at 37 °C.

### Antifungal susceptibility

The minimal inhibitory concentration (MIC) was determined by conducting broth microdilution assays adapted from the Clinical and Laboratory Standards Institute (CLSI) standard method M27-A3 (24). Serial dilutions of **5** and **8** were prepared in DMSO, resulting in a final concentration ranging from 2500 μM to 2.4 μM. Growth in RPMI-1640 medium or SC medium pH 5.5, in the case of *S. cerevisiae*, which shows poor growth in RPMI, was recorded after 24 hours of incubation at 37 °C (*C. glabrata*) or 30 °C (other *Candida spp.* and *S. cerevisiae*) by measuring OD_600_ using an Epoch Microplate Spectrophotometer and BioTek Gen5 Data Analysis Software. Growth without drug but with 2.5% DMSO (control condition) was used as the normalization condition, after background subtraction (RPMI-1640 medium). Three independent assays were performed. The MIC was defined as the concentration that inhibits at least 50% of cell growth.

### Quantification of intracellular nickel levels

*C. glabrata* cells were grown to the exponential phase and left untreated or treated overnight with the indicated concentrations of **5** or **8**, 52.2 μM of fluconazole, or the combination fluconazole and **5**, overnight. After treatment, cells were harvested and washed with 10 mM of EDTA and metal-free water. Total nickel intracellular contents were measured by Inductively Coupled Plasma Atomic Emission Spectroscopy (ICP-AES) at REQUIMTE – LAQV, Universidade Nova de Lisboa, Caparica, Portugal, as previously detailed (28). Data was normalized against OD_600._. All assays were performed with four biological replicates.

### *In vivo* fungal burden

The *Galleria mellonella* caterpillar infection model was used for *in vivo* studies, following the assay described in (61) with minor modifications. *G. mellonella* larvae were reared on a diet of pollen grains and bee wax, maintained at 25 °C in the dark and used in a final stage of development (weight of approximately 200 mg). *C. glabrata* cultures were harvested by centrifugation and suspended in PBS. Each treatment and control group included ten larvae per replicate, with three independent replicates performed, totaling 30 larvae per condition. Larvae were sequentially injected, via the hindmost left proleg sanitized with 70% (v/v) ethanol, with 5 μL of *C. glabrata* cells at a concentration of 1 x 10^9^ cells/mL, followed by a second injection with 5 μL of PBS (Control), 5 μL of compound **5** at 5.6 mM (corresponding to 312.5 μM in the hemolymph, based on hemolymph volume of approximately 90 μL) or 5 μL of compound **8** at 5.6 mM. The injections were administered at 30-minute intervals. The larvae were placed in Petri dishesin and incubated in the dark at 37 °C for 24 hours. After incubation, the larvae were punctured in the abdomen with a sterile needle and 10 μL of the hemolymph was collected and plated onto YPD agar plates supplemented with 100 μg/mL chloramphenicol for CFU counting.

### Cytotoxicity

All experiments followed the ATCC Animal Cell Culture Guidelines (62). The animal cell line HeLa was cultured in Dulbecco’s modified Eagle’s medium (DMEM, Biowest) supplemented with 10% fetal bovine serum (FBS, Sigma-Aldrich) at 37 °C in 5% CO_2_ atmosphere. For cytotoxicity experiments, HeLa cells (5 x 10^3^ cells/well) were seeded in 96-well plates. After 24 hours, different concentrations of compound **5** or compound **8** diluted in DMEM medium were added, and the plates were incubated for an additional 24 hours. Cell viability was determined using 3-(4,5-dimethylthiazol-2-yl)-2,5-diphenyltetrazolium bromide (MTT) method (63). Culture medium was removed, MTT (0.5 mg/mL in DMEM) was added to each well and incubated for 2 hours at 37 °C. Formazan crystals were solubilized with DMSO for 15 minutes and quantified spectrophotometrically at 570 nm using the Epoch Microplate Spectrophotometer and BioTek Gen5 Data Analysis Software. HeLa cell viability was expressed as the percentage of treated cells relative to the control (cells cultured in DMEM with 0.5% DMSO).

### Checkerboard assays

Assays were performed according to the CLSI standard method M27-A3 for yeast, with minor modifications (24). Stock solutions of compound **5** and fluconazole were prepared in DMSO and ultrapure water, respectively. Two-fold serial dilutions were prepared for each compound, resulting in a final concentration ranging from 0.6 to 625 µM for compound **5**, and 6.5 to 417.9 µM for fluconazole. Growth in RPMI-1640 medium was recorded after 24 h at 37 °C by measuring OD_600_ using the Epoch Microplate Spectrophotometer and Biotek Gen5 Data Analysis Software and by spotting 5 μL of each well culture onto YPD agar plates.

### Biofilm assays

The effect of **5** alone or combined with fluconazole on *C. glabrata* biofilm was assessed as previously described (64) with minor modifications. Briefly, 100 µL of exponentially growing cells diluted to a concentration of 1 x 10^7^ cells/mL was added to each well of a 96-well plate and incubated for 90 minutes at 37 °C to allow adherence. Non-adherent cells were aspirated, the wells were washed with PBS, and RPMI-1640, containing 78.1 μM of compound 5, 52.2 μM of fluconazole or the combination of both, was added. The plate was incubated for an additional 24 h at 37 °C and the inhibitory effect of the compound was evaluated by staining the biofilms with 0.4% of crystal violet (65).

### ROS measurements

Cells were grown to exponential phase and left untreated or treated with 78.1 μM of compound **5** or with 10 mM hydrogen peroxide for 2 hours. Cultures were harvested, and the OD_600_ was normalized to the lowest value. Cells were resuspended in PBS containing 5 μg/mL of dihydrorhodamine 123 (DHR123) and incubated at room temperature in the dark for 2 hours. After incubation, cells were washed three times with PBS and transferred to a black NUNC microplate for fluorescence measurement (Ex: 498 nm, Em: 539 nm) using theBioTek Plate Reader Neo2 Multimode Reader.

## Acknowledgements

This work was supported by FCT – Fundação para a Ciência e a Tecnologia, I.P., through MOSTMICRO-ITQB R&D Unit (doi.org/10.54499/UID/04612/2025, UID/PRR/4612/2025), Green-it Bioresources for Sustainability R&D Unit (UID/04551/2025, DOI: 10.54499/UID/04551/2025; UID/PRR/04551/2025, DOI: 10.54499/UID/PRR/04551/2025) and LS4FUTURE Associated Laboratory (doi.org/10.54499/LA/P/0087/2020). CM-L was a recipient of a PhD fellowship supported by FCT, with reference UI/BD/153387/2022 (doi.org/10.54499/UI/BD/153387/2022).

Cláudia Malta-Luís performed most of the experimental work, analyzed the data and wrote the manuscript draft. Giulia Romeo and Giulia Francescato designed and synthesized the complexes. Carolina Mariano contributed to preliminary experimental work. Dalila Mil-Homens performed the *in vivo* assays. Ana Petronilho provided resources and facilities, designed the compounds and supervised the chemistry part of work. Catarina Pimentel provided resources and facilities, conceptualized the project, designed the experimental approach, supervised the work and wrote the manuscript draft. All authors reviewed the manuscript and approved its final version.

## Supplementary Material

**Figure S1.** ^1^H NMR spectrum of complex **8** in DMSO-*d_6_*.

**Figure S2.** ^13^C{^1^H} NMR spectrum of complex **8** in DMSO-*d_6_*.

**Figure S3. Higher intracellular levels of nickel are observed after treatment with compound 5 than with compound 8.** The intracellular nickel content of C. glabrata cells, either untreated (Control) or treated with 2.4 μM of each compound, was quantified by ICP-AES. Data represent the mean ± standard deviation of four biological replicates. Statistical significance was assessed using one-way ANOVA with Turkey’s HSD post hoc test (*p<0.05, ****p<0.0001).

**Figure S4. Compound 5 does not induce reactive oxygen species.** Intracellular ROS levels in *C. glabrata* cells, untreated (Control) or treated with 10 mM hydrogen peroxide (H_2_O_2_) or 78.1 μM compound 5 (5), were quantified using the fluorescence dye dihydrorhodamine 123 (DHR 123). Statistical significance was assessed using one-way ANOVA with Turkey’s HSD post hoc test (***p<0.001).

## References

1. Pappas PG, Lionakis MS, Arendrup MC, Ostrosky-Zeichner L, Kullberg BJ. 2018. Invasive candidiasis. Nat Rev Dis Primers 4:18026.

2. Fisher MC, Alastruey-Izquierdo A, Berman J, Bicanic T, Bignell EM, Bowyer P, Bromley M, Bruggemann R, Garber G, Cornely OA, Gurr SJ, Harrison TS, Kuijper E, Rhodes J, Sheppard DC, Warris A, White PL, Xu J, Zwaan B, Verweij PE. 2022. Tackling the emerging threat of antifungal resistance to human health. Nat Rev Microbiol 20:557–571.

3. Campoy S, Adrio JL. 2017. Antifungals. Biochem Pharmacol 133:86–96.

4. Hoenigl M, Sprute R, Egger M, Arastehfar A, Cornely OA, Krause R, Lass-Florl C, Prattes J, Spec A, Thompson GR, 3rd, Wiederhold N, Jenks JD. 2021. The Antifungal Pipeline: Fosmanogepix, Ibrexafungerp, Olorofim, Opelconazole, and Rezafungin. Drugs 81:1703–1729.

5. WHO WHO. 2022. WHO fungal priority pathogens list to guide research, development and public health action.

6. Denning DW. 2024. Renaming *Candida glabrata*-A case of taxonomic purity over clinical and public health pragmatism. PLoS Pathog 20:e1012055.

7. Katsipoulaki M, Strappers MHT, Malavia-Jones D, Brunke S, Hube B, Gow NAR. 2024. *Candida albicans* and *Candida glabrata*: global priority pathogens. Microbiology and Molecular Biology Reviews 88.

8. Whaley SG, Berkow EL, Rybak JM, Nishimoto AT, Barker KS, Rogers PD. 2016. Azole Antifungal Resistance in *Candida albicans* and Emerging Non-*albicans Candida* Species. Front Microbiol 7:2173.

9. Cornely OA, Sprute R, Bassetti M, Chen SC, Groll AH, Kurzai O, Lass-Florl C, Ostrosky-Zeichner L, Rautemaa-Richardson R, Revathi G, Santolaya ME, White PL, Alastruey-Izquierdo A, Arendrup MC, Baddley J, Barac A, Ben-Ami R, Brink AJ, Grothe JH, Guinea J, Hagen F, Hochhegger B, Hoenigl M, Husain S, Jabeen K, Jensen HE, Kanj SS, Koehler P, Lehrnbecher T, Lewis RE, Meis JF, Nguyen MH, Pana ZD, Rath PM, Reinhold I, Seidel D, Takazono T, Vinh DC, Zhang SX, Afeltra J, Al-Hatmi AMS, Arastehfar A, Arikan-Akdagli S, Bongomin F, Carlesse F, Chayakulkeeree M, Chai LYA, Chamani-Tabriz L, Chiller T, Chowdhary A, et al. 2025. Global guideline for the diagnosis and management of candidiasis: an initiative of the ECMM in cooperation with ISHAM and ASM. Lancet Infect Dis 25:e280–e293.

10. Duggan S, Usher J. 2023. *Candida glabrata*: A powerhouse of resistance. PLoS Pathog 19:e1011651.

11. Coste AT, Kritikos A, Li J, Khanna N, Goldenberger D, Garzoni C, Zehnder C, Boggian K, Neofytos D, Riat A, Bachmann D, Sanglard D, Lamoth F, Fungal Infection Network of S. 2020. Emerging echinocandin-resistant Candida albicans and glabrata in Switzerland. Infection 48:761–766.

12. Lin Y, Betts H, Keller S, Cariou K, Gasser G. 2021. Recent developments of metal-based compounds against fungal pathogens. Chem Soc Rev 50:10346–10402.

13. Frei A, Elliott AG, Kan A, Dinh H, Brase S, Bruce AE, Bruce MR, Chen F, Humaidy D, Jung N, King AP, Lye PG, Maliszewska HK, Mansour AM, Matiadis D, Munoz MP, Pai TY, Pokhrel S, Sadler PJ, Sagnou M, Taylor M, Wilson JJ, Woods D, Zuegg J, Meyer W, Cain AK, Cooper MA, Blaskovich MAT. 2022. Metal Complexes as Antifungals? From a Crowd-Sourced Compound Library to the First In Vivo Experiments. JACS Au 2:2277–2294.

14. Cardoso J, Pinto E, Sousa E, Resende D. 2025. Redefining Antifungal Treatment: The Innovation of Metal-Based Compounds. J Med Chem 68:5006–5023.

15. Budagumpi S, Narayana BK, Keri RS, Hanumantharayudu ND, Vijayakumar M, Viswanathamurthi P. 2023. Recent advances in catalytic and electrocatalytic applications of half-sandwich nickel(0/II) N–heterocyclic carbene complexes. Applied Organometallic Chemistry 37.

16. Pawlak M, Drzezdzon J, Jarzembska KN, Jacewicz D. 2025. Optimizing Nickel(II) Complex Catalysts for High-Yield Oligomerization of Cyclohexyl Isocyanide. Inorg Chem 64:4844–4853.

17. AlAli A, Alkanad M, Alkanad K, Venkatappa A, Sirawase N, Warad I, Khanum SA. 2025. A comprehensive review on anti-inflammatory, antibacterial, anticancer and antifungal properties of several bivalent transition metal complexes. Bioorganic Chemistry 160.

18. Hopkinson MN, Richter C, Schedler M, Glorius F. 2014. An overview of N-heterocyclic carbenes. Nature 510:485–96.

19. Cazin CSJ. 2010. N-Heterocyclic Carbenes in Transition Metal Catalysis and Organocatalysis, 1 ed 10.1007/978-90-481-2866-2. Springer Dordrecht.

20. Patil SA, Patil SA, Patil R, Keri RS, Budagumpi S, Balakrishna GR, Tacke M. 2015. N-heterocyclic carbene metal complexes as bio-organometallic antimicrobial and anticancer drugs. Future Medicinal Chemistry 7:1305–1333.

21. Francescato G, Silva SMd, Leitão MIPS, Gaspar-Cordeiro A, Giannopoulos N, Gomes CSB, Pimentel C, Petronilho A. 2022. Nickel N-heterocyclic carbene complexes based on xanthines: Synthesis and antifungal activity on *Candida* sp. Applied Organometallic Chemistry 38.

22. Luca OR, Huang DL, Takase MK, Crabtree RH. 2013. Redox-active cyclopentadienyl Ni complexes with quinoid N-heterocyclic carbene ligands for the electrocatalytic hydrogen release from chemical fuels. New Journal of Chemistry 37.

23. Banach Ł, Guńka PA, Zachara J, Buchowicz W. 2019. Half-sandwich Ni(II) complexes [Ni(Cp)(X)(NHC)]: From an underestimated discovery to a new chapter in organonickel chemistry. Coordination Chemistry Reviews 389:19–58.

24. CLSI. 2017. Reference method for broth dilution antifungal susceptibility testing of yeasts. Clinical and Laboratory Standards Institute (CLSI) standard M27., 4th ed. ed.

25. Jacobs SE, Chaturvedi V. 2024. CAF to the Rescue! Potential and Challenges of Combination Antifungal Therapy for Reducing Morbidity and Mortality in Hospitalized Patients With Serious Fungal Infections. Open Forum Infect Dis 11:ofae646.

26. Cui J, Ren B, Tong Y, Dai H, Zhang L. 2015. Synergistic combinations of antifungals and anti-virulence agents to fight against *Candida albicans*. Virulence 6:362–71.

27. Van Rhijn N, White PL. 2025. Antifungal treatment strategies and their impact on resistance development in clinical settings. J Antimicrob Chemother 80:3208–3226.

28. Gaspar-Cordeiro A, Amaral C, Pobre V, Antunes W, Petronilho A, Paixao P, Matos AP, Pimentel C. 2022. Copper Acts Synergistically With Fluconazole in *Candida glabrata* by Compromising Drug Efflux, Sterol Metabolism, and Zinc Homeostasis. Front Microbiol 13:920574.

29. Kaur J, Nobile CJ. 2023. Antifungal drug-resistance mechanisms in *Candida* biofilms. Curr Opin Microbiol 71:102237.

30. Raut JS, Chauhan NM, Shinde RB, Karuppayil SM. 2013. Inhibition of planktonic and biofilm growth of *Candida albicans* reveals novel antifungal activity of caffeine. Journal of Medicinal Plants Research 7:777–782.

31. Arastehfar A, Daneshnia F, Hovhannisyan H, Fuentes D, Cabrera N, Quinteros C, Ilkit M, Ünal N, Hilmioğlu-Polat S, Jabeen K, Zaka S, Desai JV, Lass-Flörl C, Shor E, Gabaldon T, Perlin DS. 2023. Overlooked *Candida glabrata* petites are echinocandin tolerant, induce host inflammatory responses, and display poor in vivo fitness. mBio 14.

32. Brun S, Dalle F, Saulnier P, Renier G, Bonnin A, Chabasse D, Bouchara JP. 2005. Biological consequences of petite mutations in *Candida glabrata*. J Antimicrob Chemother 56:307–14.

33. Day M. 2013. Yeast Petites and Small Colony Variants: For Everything There Is a Season, p 1-41. *In* Gadd GM, Sariaslani S (ed), Advances in Applied Microbiology, vol 85. Academic Press.

34. Brun S, Berges T, Poupard P, Vauzelle-Moreau C, Renier G, Chabasse D, Bouchara JP. 2004. Mechanisms of azole resistance in petite mutants of *Candida glabrata*. Antimicrob Agents Chemother 48:1788–96.

35. Perfect JR. 2017. The antifungal pipeline: a reality check. Nat Rev Drug Discov 16:603–616.

36. Jayachandiran K, Esha S, Savitha Lakshmi M, Mahalakshmi S, Arockiasamy S. 2025. Synthesis and structural insights of bis(2-methoxy-6-[(2-methylpropyl)imino]methylphenolato) nickel (II) complex through DFT and docking investigations. Sci Rep 15:1751.

37. Benny A, Sakthivel E, M SL, Sahasranaman M, Sebastian A. 2025. Investigation of a Homoleptic Nickel(II) Complex: Synthesis, Crystal Structure, Computational Insights, and Antimicrobial Efficacy. ACS Omega 10:58369–58382.

38. Alomar K, Landreau A, Allain M, Bouet G, Larcher G. 2013. Synthesis, structure and antifungal activity of thiophene-2,3-dicarboxaldehyde bis(thiosemicarbazone) and nickel(II), copper(II) and cadmium(II) complexes: unsymmetrical coordination mode of nickel complex. J Inorg Biochem 126:76–83.

39. Malik MA, Lone SA, Wani MY, Talukdar MIA, Dar OA, Ahmad A, Hashmi AA. 2020. S-benzyldithiocarbazate imine coordinated metal complexes kill *Candida albicans* by causing cellular apoptosis and necrosis. Bioorg Chem 98:103771.

40. Ali I, Wani WA, Khan A, Haque A, Ahmad A, Saleem K, Manzoor N. 2012. Synthesis and synergistic antifungal activities of a pyrazoline based ligand and its copper(II) and nickel(II) complexes with conventional antifungals. Microb Pathog 53:66–73.

41. Liu X, Testa B, Fahr A. 2011. Lipophilicity and its relationship with passive drug permeation. Pharm Res 28:962–77.

42. Duong HP, Melechov D, Lim W, Ma J, Scroggie KR, Rajendra L, Perry B, Cruz LR, Saleem RSZ, Rutledge PJ, Motion A, van de Sande WWJ, Todd MH. 2025. Structure-activity relationships of fenarimol analogues with potent in vitro and in vivo activity against *Madurella mycetomatis*, the main causative agent of mycetoma. RSC Med Chem 16:6094–6108.

43. Roetzer A, Gabaldon T, Schuller C. 2011. From *Saccharomyces cerevisiae* to *Candida glabrata* in a few easy steps: important adaptations for an opportunistic pathogen. FEMS Microbiol Lett 314:1–9.

44. Rodrigues CF, Silva S, Henriques M. 2014. *Candida glabrata*: a review of its features and resistance. Eur J Clin Microbiol Infect Dis 33:673–88.

45. Doudican NA, Song B, Shadel GS, Doetsch PW. 2005. Oxidative DNA damage causes mitochondrial genomic instability in S*accharomyces cerevisiae*. Mol Cell Biol 25:5196–204.

46. Zhang JJ, Muenzner JK, Abu El Maaty MA, Karge B, Schobert R, Wolfl S, Ott I. 2016. A multi-target caffeine derived rhodium(i) N-heterocyclic carbene complex: evaluation of the mechanism of action. Dalton Trans 45:13161–8.

47. Dabiri Y, Schmid A, Theobald J, Blagojevic B, Streciwilk W, Ott I, Wolfl S, Cheng X. 2018. A Ruthenium(II) N-Heterocyclic Carbene (NHC) Complex with Naphthalimide Ligand Triggers Apoptosis in Colorectal Cancer Cells via Activating the ROS-p38 MAPK Pathway. Int J Mol Sci 19.

48. Truong D, Sullivan MP, Tong KKH, Steel TR, Prause A, Lovett JH, Andersen JW, Jamieson SMF, Harris HH, Ott I, Weekley CM, Hummitzsch K, Sohnel T, Hanif M, Metzler-Nolte N, Goldstone DC, Hartinger CG. 2020. Potent Inhibition of Thioredoxin Reductase by the Rh Derivatives of Anticancer M(arene/Cp*)(NHC)Cl(2) Complexes. Inorg Chem 59:3281–3289.

49. Contamine Vr, Picard M. 2000. Maintenance and Integrity of the Mitochondrial Genome: a Plethora of Nuclear Genes in the Budding Yeast. Microbiology and Molecular Biology Reviews 64:281–315.

50. Wolf K, Kaudewitz F. 1976. Effect of Caffeine on the rho^-^-induction with Ethidium Bromide in *Saccharomyces cerevisiae*. Molecular and General Genetics 146:89–93.

51. Moresi NG, Geck RC, Skophammer R, Godin D, yEvo S, Taylor MB, Dunham MJ. 2023. Caffeine-tolerant mutations selected through an at-home yeast experimental evolution teaching lab. microPublication Biology doi:10.17912/micropub.biology.000749.

52. Kan L-s, Borer PN, Cheng DM, Ts’o POP. 1980. ^1^H- and ^13^C-nmr studies on caffeine and its interaction with nucleic acids. Biopolymers 19:1641–1654.

53. Defontaine A, Bouchara JP, Declerk P, Planchenault C, Chabasse D, Hallet JN. 1999. In-vitro resistance to azoles associated with mitochondrial DNA deficiency in *Candida glabrata*. Journal of medical microbiology 48:663–670.

54. Sanglard D, Ischer F, Bille J. 2001. Role of ATP-binding-cassette transporter genes in high-frequency acquisition of resistance to azole antifungals in *Candida glabrata*. Antimicrob Agents Chemother 45:1174–83.

55. Hallstrom TC, Moye-Rowley WS. 2000. Multiple signals from dysfunctional mitochondria activate the pleiotropic drug resistance pathway in *Saccharomyces cerevisiae*. J Biol Chem 275:37347–56.

56. Kaur R, Castano I, Cormack BP. 2004. Functional genomic analysis of fluconazole susceptibility in the pathogenic yeast *Candida glabrata*: roles of calcium signaling and mitochondria. Antimicrob Agents Chemother 48:1600–13.

57. Moura TA, Junior RLR, Rocha MS. 2021. Caffeine modulates the intercalation of drugs on DNA: A study at the single molecule level. Biophys Chem 277:106653.

58. Hixon SC, Yielding KL. 1976. A Protective Effect of Caffeine on the Ethidium Bromide Induced Petite Mutation in Yeast. Mutation Research 34:195–200.

59. Ferrari S, Sanguinetti M, De Bernardis F, Torelli R, Posteraro B, Vandeputte P, Sanglard D. 2011. Loss of mitochondrial functions associated with azole resistance in *Candida glabrata* results in enhanced virulence in mice. Antimicrob Agents Chemother 55:1852–60.

60. Bouchara JP, Zouhair R, LE Boudouil S, Renier G, Filmon R, Chabasse D, Hallet JN, Defontaine A. 2000. *In-vivo* selection of an azole-resistant petite mutant of *Candida glabrata*. Journal of medical microbiology 49:977–984.

61. Pedras A, Malta-Luis C, Lima LMP, Mil-Homens D, Amaral C, Duarte AG, Antunes W, Gaspar-Cordeiro A, Louro RO, Lamosa P, Soares CM, Lousa D, Pimentel C. 2025. Caspofungin binding to iron compromises its antifungal efficacy against *Candida albicans*. Communications Biology 8:1438.

62. ATCC. Animal Cell Culture Guide. https://www.atccorg/resources/culture-guides/animal-cell-culture-guide#Glossary.

63. Ghasemi M, Turnbull T, Sebastian S, Kempson I. 2021. The MTT Assay: Utility, Limitations, Pitfalls, and Interpretation in Bulk and Single-Cell Analysis. Int J Mol Sci 22.

64. Gupta P, Gupta S, Sharma M, Kumar N, Pruthi V, Poluri KM. 2018. Effectiveness of Phytoactive Molecules on Transcriptional Expression, Biofilm Matrix, and Cell Wall Components of Candida glabrata and Its Clinical Isolates. ACS Omega 3:12201–12214.

65. Jin Y, Yip HK, Samaranayake YH, Yau JY, Samaranayake LP. 2003. Biofilm-forming ability of *Candida albicans* is unlikely to contribute to high levels of oral yeast carriage in cases of human immunodeficiency virus infection. J Clin Microbiol 41:2961–7.

