## Supplementary Figures for "A Nickel N-Heterocyclic Biscarbene Complex Derived from Caffeine Enhances Fluconazole Efficacy against *Candida glabrata*"

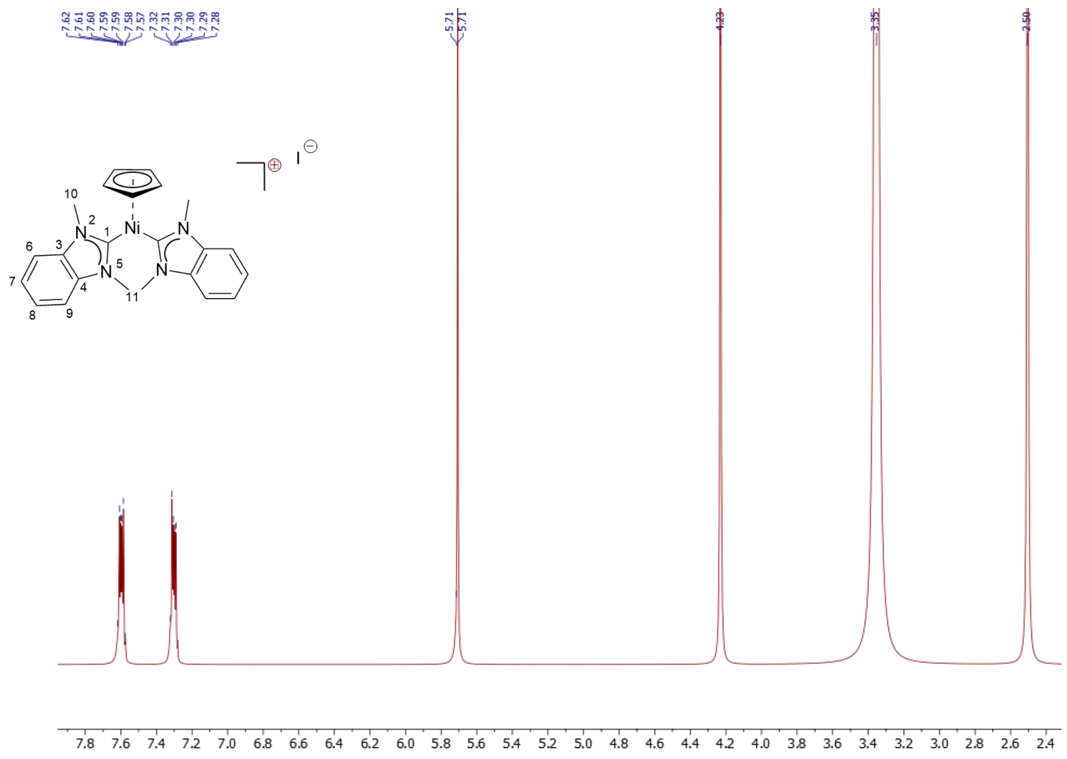


**Figure S1.** $^{1}H$ NMR spectrum of complex **8** in DMSO-$d_6$.

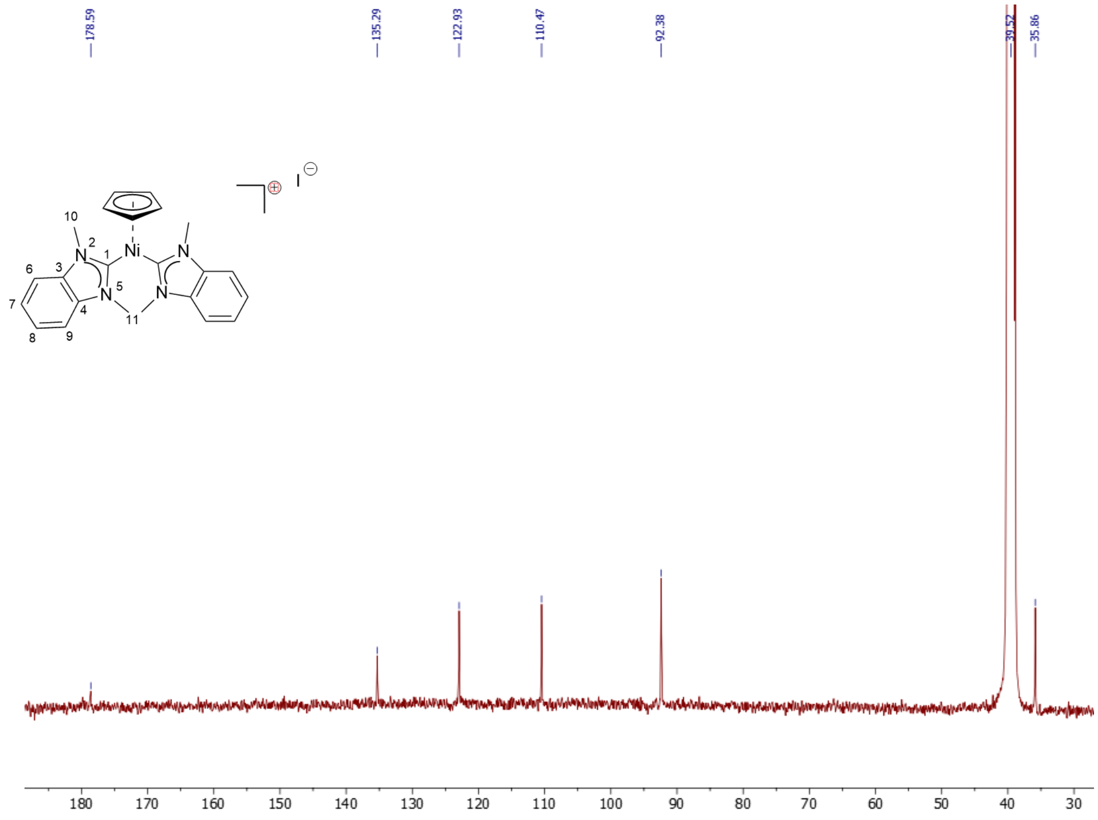


**Figure S2.** $^{13}C\{^{1}H\}$ NMR spectrum of complex **8** in DMSO-$d_6$.

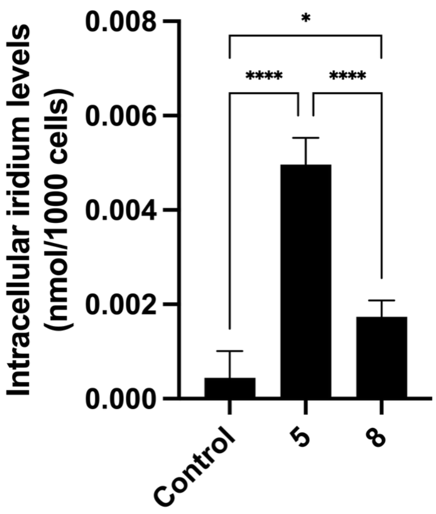


**Figure S3. Higher intracellular levels of nickel are observed after treatment with compound 5 than with compound 8.** The intracellular nickel content of C. glabrata cells, either untreated (Control) or treated with 2.4 μM of each compound, was quantified by ICP-AES. Data represent the mean ± standard deviation of four biological replicates. Statistical significance was assessed using one-way ANOVA with Turkey's HSD post hoc test (*p<0.05, ****p<0.0001).

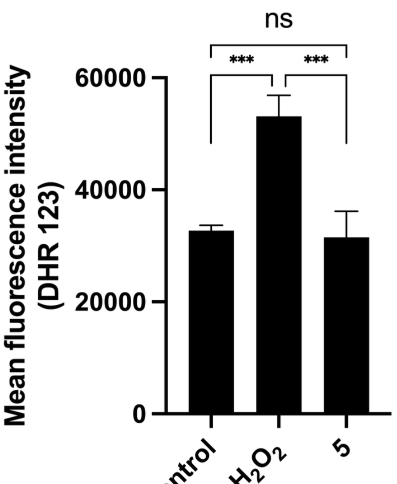


**Figure S4. Compound 5 does not induce reactive oxygen species.** Intracellular ROS levels in *C. glabrata* cells, untreated (Control) or treated with 10 mM hydrogen peroxide ($H_2O_2$) or 78.1 μM compound 5 (5), were quantified using the fluorescence dye dihydrorhodamine 123 (DHR 123). Statistical significance was assessed using one-way ANOVA with Turkey's HSD post hoc test (***p<0.001).
